# A broad and potent neutralization epitope in SARS-related coronaviruses

**DOI:** 10.1101/2022.03.13.484037

**Authors:** Meng Yuan, Xueyong Zhu, Wan-ting He, Panpan Zhou, Chengzi I. Kaku, Tazio Capozzola, Connie Y. Zhu, Xinye Yu, Hejun Liu, Wenli Yu, Yuanzi Hua, Henry Tien, Linghang Peng, Ge Song, Christopher A. Cottrell, William R. Schief, David Nemazee, Laura M. Walker, Raiees Andrabi, Dennis R. Burton, Ian A. Wilson

## Abstract

Many neutralizing antibodies (nAbs) elicited to ancestral SARS-CoV-2 through natural infection and vaccination generally have reduced effectiveness to SARS-CoV-2 variants. Here we show therapeutic antibody ADG20 is able to neutralize all SARS-CoV-2 variants of concern (VOCs) including Omicron (B.1.1.529) as well as other SARS-related coronaviruses. We delineate the structural basis of this relatively escape-resistant epitope that extends from one end of the receptor binding site (RBS) into the highly conserved CR3022 site. ADG20 can then benefit from high potency through direct competition with ACE2 in the more variable RBS and interaction with the more highly conserved CR3022 site. Importantly, antibodies that are able to target this site generally neutralize all VOCs, albeit with reduced potency against Omicron. Thus, this highly conserved and vulnerable site can be exploited for design of universal vaccines and therapeutic antibodies.

## Introduction

SARS-CoV-2 vaccines, based on the ancestral virus strain (*1, 2*), confer protective immunity and greatly decrease incidence of infection, disease severity and hospitalization from COVID-19. Many SARS-CoV-2 variants have emerged, and the designated variants of concern (VOCs), especially the recent B.1.1.529 (Omicron) variant, as well as some variants of interest, are much more resistant to neutralizing antibody responses induced by current vaccines (*3-8*). A vaccine that is highly protective against current SARS-CoV-2 VOCs could potentially provide broader protection against future emerging variants and possibly other sarbecoviruses. However, neutralizing potency and breadth are often somewhat mutually exclusive; the most highly potent nAbs target the ACE2 receptor binding site (RBS) of the spike (S) protein, but most SARS-CoV-2 VOCs have mutations in the RBS that reduce nAb binding and neutralization. Broad binding antibodies, such CR3022 that bind to other epitopes on the receptor binding domain (RBD) usually lack neutralization potency (*9*). Here, we identified a site of vulnerability on the receptor binding domain (RBD) of the SARS-CoV-2 S protein that is targeted by a few diverse antibodies. Importantly, such antibodies as exemplified by ADG20, compete with receptor binding, exhibit high neutralizing potency, and show broad activity to all known VOCs, including Omicron binding, to a conserved region that is present also on other SARS-related coronaviruses including SARS-CoV-1, WIV1 and SHC014.

## Results

Some of the authors previously developed mAb ADG20 as an extended half-life version of potent-and-broad human antibody ADG-2 (*10, 11*). ADG20 and ADG-2 share the same antigen-binding fragment (Fab) domain with a few amino-acid changes in the fragment crystallizable region (Fc region) (*11*). ADG20 and ADG-2 neutralize a broad spectrum of SARS-related coronaviruses including SARS-CoV-2, SARS-CoV-1, WIV-1 and SHC014 with high potency (IC_50_ ranging from 1 to 30 ng/ml against authentic viruses), as well as confer outstanding protection in mice infected with SARS-CoV-1 or SARS-CoV-2 (*10*). ADG20 is now in phase II/III trials for COVID-19 treatment and prevention (*12, 13*). A low resolution cryo-electron microscopy (cryo-EM) structure (∼6 Å) of ADG20 Fab was previously reported in complex with SARS-CoV-2 S protein (*10*).

Here, we determined a crystal structure of ADG20 Fab in complex with the wild-type SARS-CoV-2 RBD to 2.75 Å resolution to decipher the atomic details of the antibody-antigen interactions and the molecular features of this site of vulnerability (Tables S1 and S2). ADG20 targets one corner of the RBD that is distant from the ridge region (Figure 1A-C) through CDRs H1, H2, H3, L1, and L3 (Figure 1B). The buried surface areas (BSA) of SARS-CoV-2 RBD conferred by the heavy and light chains of ADG20 are 488 and 204 Å^2^, respectively. The epitope of ADG20 overlaps with the RBS (Figure 1A, B), and binding of the antibody would clash with ACE2 binding to the RBD (Figure 1C). The epitope of ADG20 is only accessible when RBD is in the “up” conformation (Figure S1). CDRs H1 and H2 of ADG20 participate in a network of interactions with the RBD (Figure 1D), where V_H_ E52a forms a hydrogen bond and salt bridge with Y505 and R403, respectively. R403 is further stabilized by D405, which hydrogen bonds with V_H_ S56 and V_H_ Y33. V_H_ Y33 in turn stacks with Y505. V_H_ Y55 interacts with R408 through a cation-π interaction (Figure 1D). CDR H3 forms five hydrogen bonds with the RBD (Figure 1E). The light chain of ADG20 is also involved in RBD recognition, where V_L_ Y91, L95, and L95c form a hydrophobic pocket to accommodate V503. V_L_ Y91 and Y31 hydrogen bond with V503 and Q506, respectively (Figure 1F). G504 is also involved in interaction with ADG20 (Figure 1G). In our escape mutation study, RBD-G504D emerged in a second passage of authentic SARS-CoV-2 in the presence of ADG20 and exhibited full escape (Figure S2), consistent with our previous finding where G504D abrogates binding of ADG-2 to the SARS-CoV-2 RBD (*10*), illustrating the importance of this interaction.

**Figure 1.**
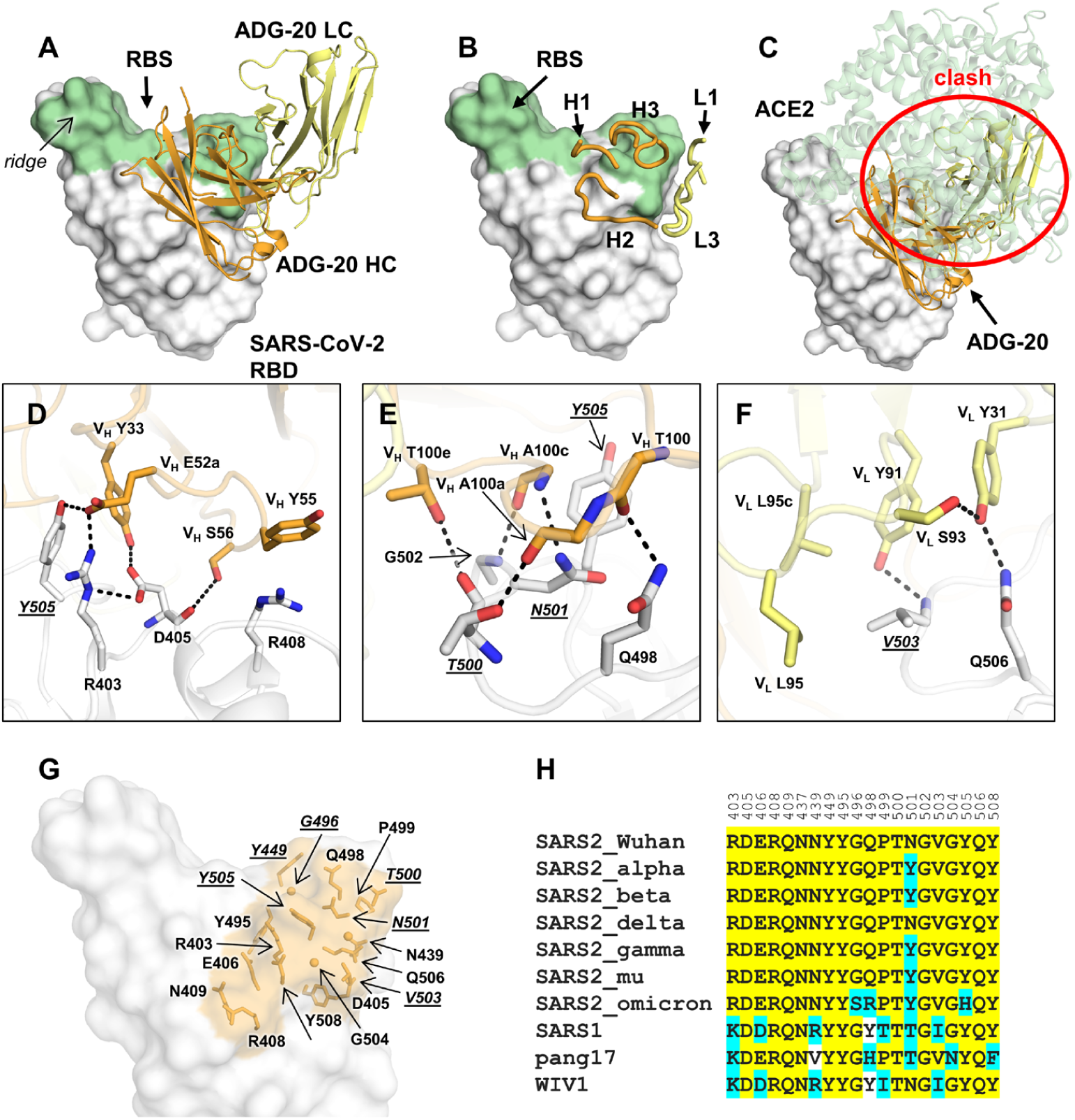
Crystal structure of ADG20 in complex with SARS-CoV-2 RBD. The heavy and light chains of ADG20 are shown in orange and yellow, respectively. SARS-CoV-2 RBD is shown in white. ADG20 epitope residues and RBS residues are defined by residues with surface area buried by ADG20 and ACE2 > 0 Å^2^, respectively, calculated based on the ADG20/RBD crystal structure in this study and an RBD/ACE2 complex structure (PDB 6M0J) (*48*) by PISA (*44*). **(A)** Structure of ADG20 Fab in complex with SARS-CoV-2 RBD. The receptor binding site (RBS) is shown in green. For clarity, the constant domains of ADG20 Fab were omitted. **(B)** ADG20 CDRs that interact with SARS-CoV-2 are shown as tubes. **(C)** The ACE2/RBD complex structure is superimposed on the ACE-2/RBD complex. ADG20 would clash with ACE2 (green) if bound to RBD simultaneously (indicated by red circle). **(D-F)** Detailed interactions between ADG20 **(D)** CDRs H1 and H2, **(E)** CDR H3, and **(F)** light chain with SARS-CoV-2 RBD. Hydrogen bonds and salt bridges are represented by black dashed lines. Epitope residues involved in ACE2 binding are underlined and italicized. Kabat numberings are used for antibody residues throughout this paper. **(G)** Epitope residues of ADG20 are highlighted in orange, with side chains shown as sticks and Cα of glycines as spheres. **(H)** Sequence alignment of the ADG20 epitope residues in a subset of SARS-like viruses where we also generated PSV neutralization data. Conserved residues among SARS-related coronaviruses are highlighted with a yellow background, while similar residues are in a cyan background [amino acids scoring greater than or equal to 0 in the BLOSUM62 alignment score matrix (*49*) were counted as similar here].

The ADG20 epitope residues are generally conserved among SARS-CoV-2 and its variants, as well as other SARS-related coronaviruses including SARS-CoV-1 (Figure 1G, H). Notably, unlike some major classes of RBD-targeting neutralizing antibodies (e.g. IGHV3-53 and IGHV1-2 antibodies), which are sensitive to mutations in Beta and Gamma variants (*14*), all of the ADG20 epitope residues are conserved among VOCs Alpha, Beta, Gamma, and Delta, except for N501Y (Figure 1H), which only minimally affects the interaction with ADG20 (Figure S3). The SARS-CoV-2 Omicron variant fully escapes neutralization by 14 out of 18 tested mAbs and cocktail pairs. By contrast, ADG20 retains neutralization activity against Omicron (IC_50_ = 1.2 μg/ml), although with approximately 100-fold reduction compared to ancestral SARS-CoV-2 (IC_50_ = 12 ng/ml) (Figure 2A). The neutralization activity of ADG20 against Omicron is comparable with the Evushield cocktail of antibodies AZD1061+AZD8895 (IC_50_ = 1.3 μg/ml), but is much more potent against SARS-CoV-1 and other sarbecoviruses (IC_50_ = 2-19 ng/ml) in the panel of viruses tested (Figure 2A). Sotrovimab (derived from S309), a clinically authorized antibody for emergency use, is less affected by the Omicron variant, but because its starting potency against WT is lower, its absolute IC_50_ (0.9 μg/ml) against Omicron is similar to ADG20. Thus, ADG20 is one of the most potent nAbs among all tested antibodies against a panel of viruses (Figure 2A). Our observations are consistent with previously reported neutralization results performed with pseudotyped and authentic viruses (*15-17*). Several ADG20-epitope contact residues from SARS-CoV-2 differ in SARS-CoV-1, pang17, and WIV1 (Figure 1H), but apparently can be accommodated by ADG20 (Figure S3).

**Figure 2.**
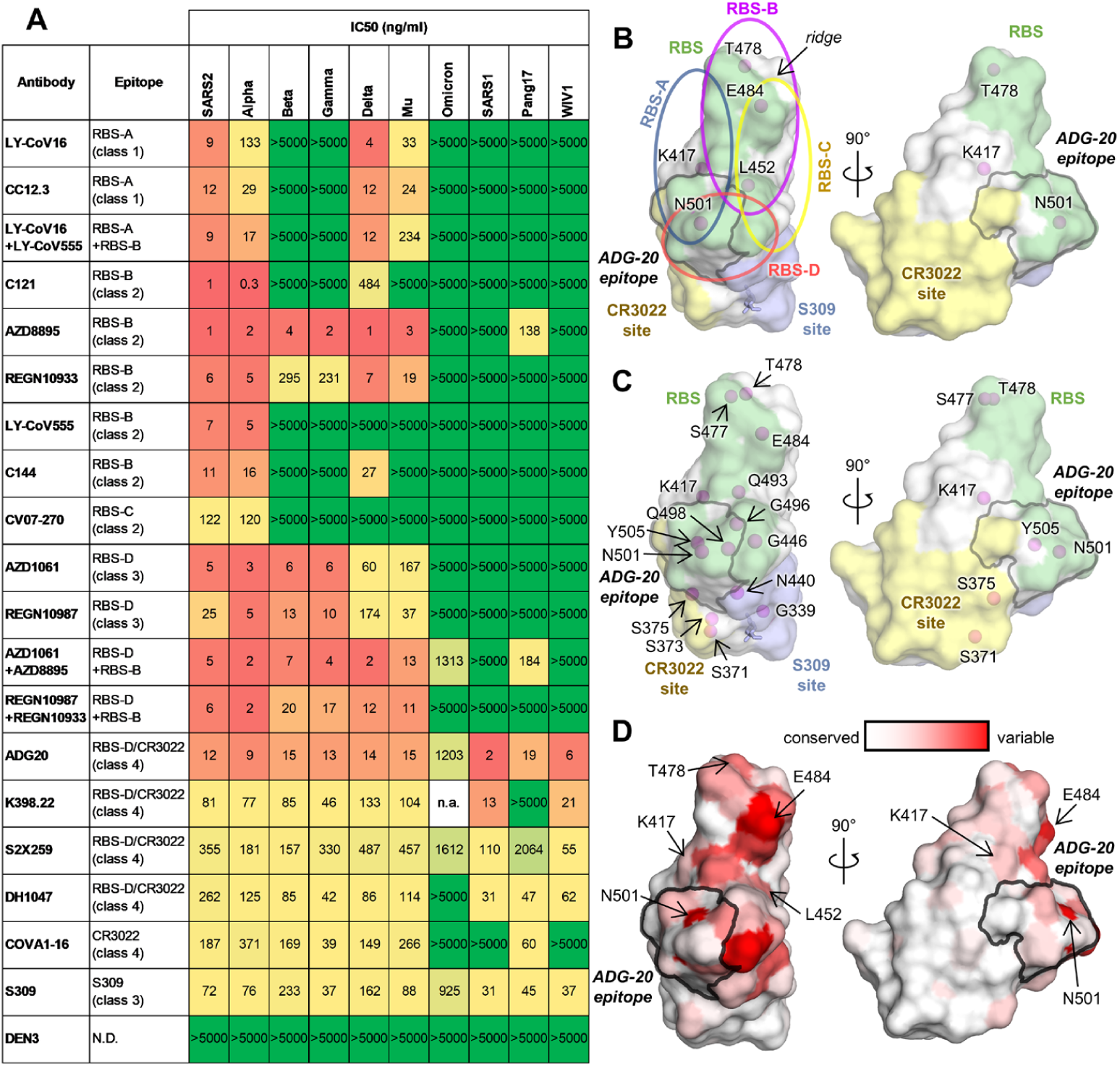
Neutralization of antibodies targeting different epitopes against SARS-CoV-2 variants and other SARS-related coronaviruses. **(A)** Neutralization of antibodies against pseudotyped SARS-CoV-2 variants and other SARS-related coronaviruses. Ancestral SARS-CoV-2 is shown as “SARS2”. Epitope classes are as defined in (*14*). Antibody epitopes that include residues in both RBS-D and CR3022 sites are shown as “RBS-D/CR3022”. “n.a.” = data not available. Corresponding categories (class 1-4) that were classified in (*50*) are shown in brackets. **(B-C)** A map of RBD epitopes and mutated residues in SARS-CoV-2 variants of concern (VOCs). The receptor binding site (RBS), CR3022 site, and S309 site are shown in green, yellow, and lavender, respectively. The ADG20 epitope is highlighted with a grey outline. Mutated residues in SARS-CoV-2 VOCs Alpha, Beta, Gamma, and Delta are labeled in (B), and those mutated in the VOC Omicron are labeled in (C). The N343 glycan is shown as sticks. Four RBS subsites, RBS-A, B, C, and D are indicated by circles in (B) but omitted in (C) for clarity. Antibodies targeting the RBS-A/class 1 epitope are mainly encoded by IGHV3-53/3-66 genes. This class of antibodies is generally sensitive to mutations K417N/T in Beta, Gamma, and Omicron (Figure S6A). Many RBS-B/class 2 [including another major class of antibodies that are encoded by IGHV1-2 and share structural convergence (Figure S6B)] and RBS-C antibodies are often sensitive to E484 mutations in the ridge region of the RBD. L452 is located on one edge of the RBS and interacts with many RBS-B and RBS-C antibodies (Figure S6C). N501 is located in the RBS-A and RBS-D epitopes, but N501Y, the only mutated RBD residue in the Alpha variant, is often tolerated by RBS-A and RBS-D antibodies (Figure S3). The Alpha variant not unexpectedly has the least immune escape among all VOCs. **(D)** Sequence variation of 14 SARS-related coronaviruses (including ancestral SARS-CoV-2 and variants Alpha, Beta, Gamma, Delta, Mu, Omicron, and SARS-CoV-1, BM4831, BtKY72, pang17, RaTG13, RsSHC014, and WIV1) mapped onto an RBD structure [from white (low) to red (high)] The ADG20 epitope is highlighted with a grey outline. Mutated residues in SARS-CoV-2 VOCs are indicated by arrows.

ADG20 is an affinity-matured progeny of ADI-55688, a broad RBD-targeting monoclonal antibody isolated from a SARS-CoV-1-convalescent donor (*10, 18*). Like ADG20, ADI-55688 cross-reacts with RBDs of SARS-CoV-2 and SARS-CoV-1 and neutralizes both viruses (*10*). ADI-55688 differs from ADG20 by only five amino acids, with three located in the heavy chain and two in the light chain (Figure S4A). These mutated residues in ADG20 confer a nearly 200-fold improved binding affinity and a 100-fold increased neutralizing activity against SARS-CoV-2 compared to ADI-55688 (*10*). We also determined a crystal structure of ADI-55688 Fab in complex with SARS-CoV-2 RBD at 2.85 Å and compared with the ADG20/RBD structure. These two antibodies target the same epitope through a near-identical binding approach (Figure S4B). The V_H_ S52a substitution by glutamic acid in ADG20 leads to a salt bridge with RBD-R403 (Figure S4C), and the V_H_ W100b mutation from valine increases interaction with V_L_ H34 and V_H_ F96. These substitutions appear to stabilize the conformation of the light and heavy chain CDRs (Figure S4C) and result in an improved off-rate (*10*).

RBD is the major target of neutralizing antibodies against SARS-CoV-2 (*19*). To understand the differential effects of binding and neutralization by ADG20 compared to other RBD antibodies, we first mapped all of the VOC mutations in Alpha, Beta, Gamma, and Delta onto the RBD epitope classes. Only five mutated residues K417, L452, T478, E484, and N501 are located in the RBD in these VOCs and are distributed throughout the RBS and cover all four RBS epitopes (Figure 2B). In contrast, 15 residues are mutated in Omicron RBD (G339D, S371L, S373P, S375F, K417N, N440K, G446S, S477N, T478K, E484A, Q493R, G496S, Q498R, N501Y, and Y505H), where 8 of these 15 residues are directly involved in ACE2 binding (Figures 2C and S5). The RBS is the most variable site among SARS-related viruses (Figure 2D), but this high variation is tolerated by the receptor. In contrast, none of the mutations of VOCs Alpha, Beta, Gamma, and Delta are found in the relatively conserved CR3022 and S309 sites, although 4 of the 15 Omicron mutations are located in either the CR3022 (371 and 375) or S309 (339 and 440) sites (Figure 2B, C). Previously, we classified the RBD-targeting antibodies into six sites: RBS sites RBS-A, B, C, and D, CR3022 site and S309 site (Figure 2B) (*14*). These antibodies are often encoded by different germline genes with different sensitivities to VOC mutations (*14*). We show here the impact of the Omicron mutations on all of these sites (Figure 2C). Mutations in the 371-375 region (S371L, S373P, and S375F) of the Omicron variant induce a backbone shift away from the ADG20 paratope (Figure S5A) and decrease ADG20 neutralization (*16*). In addition, four mutations in Omicron reside within the ADG20 epitope (Figures 1G, H, and S5B), but single mutations of these four residues minimally alter ADG20 neutralization (*16*).

To gain further insights into ADG20 protection against SARS-CoV-2 and other sarbecoviruses, we further analyzed the binding and neutralizing activities of representative antibodies targeting the different class of RBD epitopes, including some of the antibodies authorized for COVID-19 therapeutic prevention and/or treatment against SARS-CoV-2 and VOCs (Figure S3). We then compared the neutralization potency and breadth of each antibody versus ADG20 in a potency vs. breadth plot (Figure 3A). Although most RBS antibodies are potent against the ancestral SARS-CoV-2, they are sensitive to VOC mutations suggesting that potency is usually associated with a tradeoff in breadth. Most tested RBS-A, B, and C antibodies can be escaped by at least one VOC (Figure 2A) due to the mutated residues being largely located in the RBS (Figure 2B, C). In contrast, the CR3022 site is much more conserved than the RBS (Figure 2C) (*9, 14, 19*). CR3022-site antibody COVA1-16 is a broadly neutralizing antibody (*20*), but is less potent than most RBS antibodies and loses neutralization capabilities against Omicron. Other CR3022-site targeting antibodies, e.g., S304 and CR3022, have also been reported to have broad binding breadth, but low or no neutralization potency to SARS-CoV-2 (*9, 21, 22*). Binding breadth but limited neutralization potency for an antibody may seem a paradox in many cases but may be due to insufficient affinity, relative inaccessibility of the epitope on the spike, or inability to directly compete with the ACE2 receptor.

**Figure 3.**
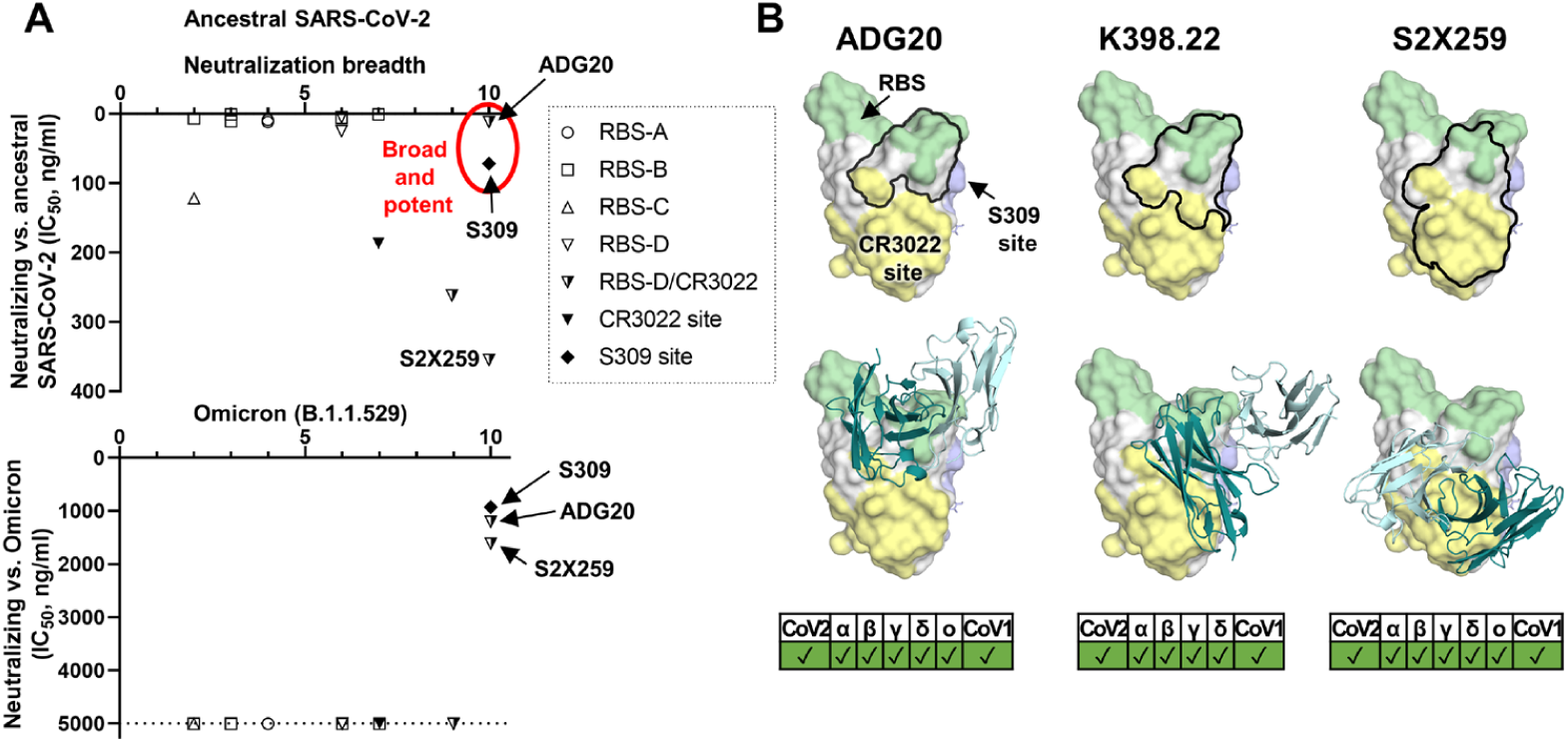
A common region on the RBD targeted by potent-and-broad neutralizing antibodies against SARS-CoV-2. **(A)** A scatter plot of antibody neutralization potency (IC_50_, ng/ml) against (top) ancestral SARS-CoV-2 and (bottom) Omicron vs. neutralization breadth, defined by the number of SARS-related coronavirus strains neutralized in this study (ten strains in total, including ancestral SARS-CoV-2, Alpha, Beta, Delta, Gamma, Mu, Omicron, SARS-CoV-1, pang17, and WIV1). **(B)** All of the potent-and-broad neutralizing antibodies target a region that spans from one end to the RBS (green) to the proximal CR3022 site (yellow). Epitopes of each antibody are outlined by black lines (top), where the variable domains of ADG20, K398.22, and S2X259 are shown in cartoon representation with heavy chains in deep teal and light chains in light cyan (bottom). Epitopes and the receptor binding site are defined by residues with BSA > 0 Å^2^ as calculated by PISA (*44*) using structures of ADG20 (this study), K398.22 (*23*), S2X259 (PDB 7RAL) (*51*), CR3022 (PDB 6W41) (*9*), S309 (PDB 6WPS) (*21*), and ACE2 (PDB 6M0J) (*48*). SARS-CoV-2 variants as well as SARS-CoV-1 can be neutralized by these antibodies, as shown at the bottom of the panel. Omicron data for K398.22 are not available.

Notwithstanding, a particularly noteworthy epitope is targeted by a few potent-and-broad antibodies including ADG20 (Figures 2A and 3). All three tested antibodies (ADG20, DH1047, and S2X259) exhibited high (<20 ng/ml) to moderate (40-500 ng/ml) potency to CoVs that include SARS-CoV-2, Alpha, Beta, Gamma, and Delta VOCs, and other SARS-related coronaviruses including SARS-CoV-1, pang17, and WIV1, with ADG20 exhibiting high potency to all of these CoVs (Figure 2A). DH1047 and ADI-55688, which was evolved in vitro to produce ADG20, were both isolated from a SARS-CoV-1 convalescent donor. This epitope seems infrequently targeted by the SARS-CoV-2 antibody repertoire, probably explaining why mAbs directed to this site are rarely isolated and described in the literature. Importantly, unlike most potently neutralizing antibodies targeting other antigenic sites on the RBD, ADG20 and S2X259 also demonstrated neutralization activity to Omicron (Figure 2A). These antibodies target a site on the RBD spanning the RBS-D and CR3022 sites (Figure 3) and are encoded by various germline genes with distinct CDR H3 sequences (Table S3). Some of us previously reported macaque antibody K398.22 that targets a similar region and also neutralizes a wide range of SARS-CoV-2 VOCs and sarbecoviruses (*23*) (Figure 3, Table S3). DH1047 also targets the RBS-D/CR3022 site and neutralizes most SARS-CoV-2 VOCs and SARS-CoV-1, but did not show neutralization activity against Omicron at the highest concentration of antibody used (5,000 ng/ml) (Figure S7). This effect may be a consequence of its relatively low neutralization against the ancestral SARS-CoV-2 (IC_50_ = 262 ng/ml) (Figure 2A) and slight differences in the key contact residues. This result is consistent with a previous study where DH1047 exhibited low neutralization activity (IC_50_ ∼ 10,000 ng/ml) against Omicron PSV (*16*). Antibodies targeting the other side of the RBS-D site (Figure S7), AZD1061 and REGN10987, are not able to neutralize the Omicron variant or other tested SARS-related coronaviruses such as SARS-CoV-1, pang17, and WIV1, possibly due to the relatively low conservation of this sub-epitope (Figures 2A, C and S7A). LY-CoV1404 also binds to the same side of RBS-D and was reported to neutralize Omicron but not bind SARS-CoV-1 (*24*). Thus, antibodies like ADG20 antibodies take advantage of the properties of two sites: targeting the RBS region that confers direct competition with receptor binding and strong potency and, on the other hand, targeting the conserved CR3022 part that imparts breadth not only against SARS-CoV-2 VOCs but also other SARS-related coronaviruses.

## Discussion

The Omicron variant is spreading globally at an unprecedented rate with the highest level of immune evasion so far in all observed variants of concern (*3-5*). Omicron is resistant, or has reduced effectiveness, to most current authorized therapeutic monoclonal antibodies as well as cocktails, including LY-CoV555, LY-CoV016, REGN10933, REGN10987, BRII-196, etc. However, neutralization by S309 or its derivative Sotrovimab/VIR-7831 was only reduced by 3-10 fold against Omicron (*5, 15, 16, 25-27*) compared to 40-100 fold reduction for ADG-2/ADG20 against Omicron pseudotyped virus here (Figure 2A) and in another study (*16*). Importantly, using authentic viruses, only a 20-fold reduction of ADG20 neutralization against Omicron was observed compared to Delta, and was the most potent among all the tested therapeutic antibodies (*15*). Thus, the S309 site targeted by Sotrovimab/VIR-7831 is another vulnerable site for broadly neutralizing antibodies. S309 exhibits broad neutralization against all SARS-CoV-2 VOCs as well as other SARS-related coronaviruses (*21, 28, 29*). However, S309-site antibodies show no or little neutralization in some PSV systems (*29, 30*), which could be due to glycoform differences. This finding raises another question of whether vaccine-elicited antigen proteins possess different glycoforms from authentic viruses. Glycan profiling of authentic virus and antigen proteins elicited by mRNA SARS-CoV-2 or other vaccines may address this question in future studies.

In summary, we identify a vulnerable site on the SARS-CoV-2 RBD that antibodies can target and mitigate against the mutations found in VOCs. These relatively rare antibodies to date extend their binding interactions from the RBS, where they can directly compete with ACE2 binding, to the most highly conserved site on the RBD where their overall footprint on the RBD confers both neutralization potency and breadth. We recently reported a class of antibodies that possess long CDRs H3 with a ‘YYDRxG’ motif that target the highly conserved CR3022 site. Such antibodies can block ACE2 without direct binding to the RBS (*31*), a few of which exhibit impressive breadth and potency to a wide range of SARS-CoV-2 VOCs and sarbecoviruses (*31*). The neutralization effectiveness of antibodies generated during infection or vaccination have been substantially reduced by emerging SARS-CoV-2 variants (*3-5, 32-34*). The VOC mutations until Omicron have been largely confined to the RBS, yet these same mutations do not adversely affect receptor binding and viral entry in host cells. Universal vaccines or antibody therapeutics that are insensitive or less susceptible to SARS-CoV-2 mutations are urgently needed to protect against the continuous antigenic drift of the virus (*35*). Several universal vaccine designs have been proposed and tested, including mosaic nanoparticles conjugated with various RBDs (*36*), chimeric spike mRNA-based vaccines (*37*), etc. Notwithstanding, such vulnerable sites in the RBD may currently be a desirable target for universal vaccine design and for antibody therapeutics.

## MATERIALS AND METHODS

### Expression and purification of IgGs and Fabs

The heavy and light chains were cloned into phCMV3. The plasmids were transiently co-transfected into ExpiCHO cells at a ratio of 2:1 (HC:LC) using ExpiFectamine™ CHO Reagent (Thermo Fisher Scientific) according to the manufacturer’s instructions. The supernatant was collected at 10 days post-transfection. The IgGs and Fabs were purified with a CaptureSelect™ CH1-XL Affinity Matrix (Thermo Fisher Scientific) followed by size exclusion chromatography.

### Crystallization and structure determination

Expression and purification of the SARS-CoV-2 spike receptor-binding domain (RBD) for crystallization were as described previously (*9*). Briefly, the RBD (residues 333-529) of the SARS-CoV-2 spike (S) protein (GenBank: QHD43416.1) was cloned into a customized pFastBac vector (*38*), and fused with an N-terminal gp67 signal peptide and C-terminal His_6_ tag (*9*). A recombinant bacmid DNA was generated using the Bac-to-Bac system (Life Technologies). Baculovirus was generated by transfecting purified bacmid DNA into Sf9 cells using FuGENE HD (Promega), and subsequently used to infect suspension cultures of High Five cells (Life Technologies) at an MOI of 5 to 10. Infected High Five cells were incubated at 28 °C with shaking at 110 r.p.m. for 72 h for protein expression. The supernatant was then concentrated using a 10 kDa MW cutoff Centramate cassette (Pall Corporation). The RBD protein was purified by Ni-NTA, followed by size exclusion chromatography, and buffer exchanged into 20 mM Tris-HCl pH 7.4 and 150 mM NaCl.

ADG-20/RBD and ADI-55688/RBD complexes were formed by mixing each of the protein components at an equimolar ratio and incubating overnight at 4°C. The protein complex was adjusted to 12 mg/ml and screened for crystallization using the 384 conditions of the JCSG Core Suite (Qiagen) on our robotic CrystalMation system (Rigaku) at Scripps Research. Crystallization trials were set-up by the vapor diffusion method in sitting drops containing 0.1 μl of protein and 0.1 μl of reservoir solution. For the ADG-20/RBD complex, optimized crystals were then grown in drops containing 0.1 M sodium citrate, pH 4.16 and 1.45 M ammonium sulfate at 20°C. Crystals appeared on day 7, were harvested on day 15 by soaking in reservoir solution supplemented with 15% (v/v) ethylene glycol, and then flash cooled and stored in liquid nitrogen until data collection. Diffraction data were collected at cryogenic temperature (100 K) at beamline 23-ID-B of the Advanced Photon Source (APS) at Argonne National Labs. For the ADI-55688/RBD complex, optimized crystals were then grown in drops containing 0.08 M sodium acetate, pH 3.8, 1.6 M ammonium sulfate, and 20% (v/v) glycerol at 20°C. Crystals appeared on day 7, were harvested on day 10 by soaking in reservoir solution supplemented with 20% (v/v) ethylene glycol, and then flash cooled and stored in liquid nitrogen until data collection. Diffraction data were collected at cryogenic temperature (100 K) at the Stanford Synchrotron Radiation Lightsource (SSRL) on Scripps/Stanford beamline 12-1. Diffraction data were processed with HKL2000 (*39*). Structures were solved by molecular replacement with PHASER (*40*) using models of the RBD and COVA2-39 derived from PBD 7JMP (*41*). Iterative model building and refinement were carried out in COOT (*42*) and PHENIX (*43*), respectively. Epitope and paratope residues, as well as their interactions, were identified by accessing PISA at the European Bioinformatics Institute (http://www.ebi.ac.uk/pdbe/prot_int/pistart.html) (*44*).

### Biolayer interferometry binding assay

RBD proteins for the biolayer interferometry (BLI) binding assay were expressed in human cells. RBDs were cloned into phCMV3 vector and fused with a C-terminal His_6_ tag. The plasmids were transiently transfected into Expi293F cells using ExpiFectamine 293 Reagent (Thermo Fisher Scientific) according to the manufacturer’s instructions. The supernatant was collected at 7 days post-transfection. The His_6_-tagged proteins were then purified with Ni Sepharose Excel protein purification resin (Cytiva) followed by size exclusion chromatography. Omicron RBD was purchased from ACROBiosystems Inc.

The BLI assays were performed by using an Octet Red instrument (FortéBio) as described previously (*9*). To measure the binding kinetics of anti-SARS-CoV-2 IgGs and RBDs, the IgGs were diluted with kinetic buffer (1x PBS, pH 7.4, 0.01% BSA and 0.002% Tween 20) into 15 µg/ml. The IgGs were then loaded onto anti-human IgG Fc (AHC) biosensors and interacted with 5-fold gradient dilution (500 nM – 20 nM) of SARS-CoV-2 RBDs, and 500 nM of RBDs of SARS-related coronaviruses. The assay consisted of the following steps. 1) baseline: 1 min with 1x kinetic buffer; 2) loading: 90 seconds with IgGs; 3) wash: 15 seconds wash of unbound IgGs with 1x kinetic buffer; 4) baseline: 1 min with 1x kinetic buffer; 5) association: 90 seconds with RBDs; and 6) dissociation: 90 seconds with 1x kinetic buffer. For estimating *K*_D_, a 1:1 binding model was used.

### Pseudovirus neutralization assay

Pseudovirus (PSV) preparation and assays were performed as previously described with minor modifications (*45*). Pseudovirions were generated by co-transfection of HEK293T cells with MLV-gag/pol (Addgene #14887) and MLV-Luciferase (Addgene #170575) plasmids and SARS-CoV-2 spike WT or variants with an 18-AA truncation at the C-terminus. Supernatants containing pseudotyped virus were collected 48 h after transfection and frozen at -80°C for long-term storage. PSV neutralizing assay was carried out as follows. 25 μl serial dilution of purified antibodies in DMEM with 10% heat-inactivated FBS, 4mM L-Glutamine and 1% P/S were incubated with 25 µl PSV at 37°C for 1 h in 96-well half-well plate (Corning, 3688). After incubation, 10,000 Hela-hACE2 cells were added to the mixture with 20 µg/ml Dextran (Sigma, 93556-1G) to enhance infectivity. At 48 h post incubation, the supernatant was aspirated, and HeLa-hACE2 cells were then lysed in luciferase lysis buffer (25 mM Glegly pH 7.8, 15 mM MgSO4, 4 mM EGTA, 1% Triton X-100). Bright-Glo (Promega, E2620) was added to the mixture following the manufacturer’s instruction, and luciferase expression was read using a luminometer. Samples were tested in duplicate, and assays were repeated at least twice for confirmation. Fifty percent maximal inhibitory concentrations (IC_50_), the concentrations required to inhibit infection by 50% compared to the controls, were calculated using the dose-response-inhibition model with 5-parameter Hill slope equation in GraphPad Prism 7 (GraphPad Software).

### SARS-CoV-2 escape assay

Escape assays in the presence of ADG20 was performed using authentic SARS-CoV-2, as previously described (*46*). Briefly, 105 TCID50 of an early, Wuhan-like SARS-CoV-2 strain (2019-nCoV/Italy/INMI1) was added to serial dilutions of ADG20 IgG ranging from 4.9 ng/mL to 10,000 ng/mL. The mixture was incubated for 1 h at 37 °C, 5% CO_2_ before adding to a 24-well plate coated in a subconfluent Vero E6 cell monolayer. The plate was incubated for 5 days at 37 °C, 5% CO_2_ and examined for signs of cytopathic effect (CPE). Viral sample at the lowest mAb dilution exhibiting complete CPE was used as the stock for the subsequent passage. Virus was also passaged in the absence of antibody to control for tissue-culture adaptations that arise independent of antibody pressure. At each passage, both the no-antibody control and virus pressured with ADG20 were harvested, propagated in 25-cm^2^ flasks, and aliquoted at -80°C for RNA extraction, RT-PCR, and sequencing.

### Next-generation sequencing of virus escape variants

NGS of the SARS-CoV-2 spike gene was performed at Science Park. Viral RNA was reverse transcribed and prepared for NGS using the Swift Amplicon SARS-CoV-2 research panel (Swift Biosciences, Ann Arbor, MI USA), following the manufacturer’s instructions. Libraries were quantified by qPCR using Kapa Lib Quant Kit (Roche Diagnostics), pooled at equimolar concentrations, and sequenced using the Illumina MiSeq system (2×250bp paired-end mode). Sequences were trimmed using Cuadapt v2.8 and consensus sequences, defined as sequence present in >50% of reads, were generated via de novo sequence construction using MegaHit (*47*).

## ACKNOWLEDGEMENTS

We thank Robyn Stanfield for assistance in data collection. This work was supported by the Bill and Melinda Gates Foundation INV-004923 (I.A.W., D.R.B.) and by a grant from Adagio (W.R.S., D.R.B., I.A.W.). We are grateful to the staff of Advanced Photon Source and Stanford Synchrotron Radiation Lightsource (SSRL) Beamline 12-1 for assistance. GM/CA@APS has been funded by the National Cancer Institute (ACB-12002) and the National Institute of General Medical Sciences (AGM-12006, P30GM138396). This research used resources of the Advanced Photon Source; a U.S. Department of Energy (DOE) Office of Science User Facility operated for the DOE Office of Science by Argonne National Laboratory under Contract No. DE-AC02-06CH11357. Extraordinary facility operations were supported in part by the DOE Office of Science through the National Virtual Biotechnology Laboratory, a consortium of DOE national laboratories focused on the response to COVID-19, with funding provided by the Coronavirus CARES Act. Use of the Stanford Synchrotron Radiation Lightsource, SLAC National Accelerator Laboratory, is supported by the U.S. Department of Energy, Office of Science, Office of Basic Energy Sciences under Contract No. DE-AC02-76SF00515. The SSRL Structural Molecular Biology Program is supported by the DOE Office of Biological and Environmental Research, and by the National Institutes of Health, National Institute of General Medical Sciences (P30GM133894). The contents of this publication are solely the responsibility of the authors and do not necessarily represent the official views of NIGMS or NIH.

## AUTHOR CONTRIBUTIONS

M.Y., X.Z., and I.A.W. designed the study. M.Y., X.Z., C.Y.Z., X.Y., H.L., W.Y., Y.H., C.A.C., and W.R.S. conducted protein construction, expression, and purification. M.Y., X.Z. and H.T. performed the crystallization, M.Y. and X.Z. collected X-ray data, determined, and refined the X-ray structures. W.H., P.Z., T.C., L.P., G.S., performed the neutralization assays under supervision of D.N., R.A., and D.R.B. M.Y. carried out the binding assays. C.I.K. and L.M.W. provided the ADG20 antibody and carried out viral escape experiments. M.Y., X.Z. and I.A.W. wrote the paper and all authors reviewed and/or edited the paper.

## COMPETING INTERESTS

C.I.K. and L.M.W. are employees of Adimab, LLC and hold shares in Adimab, LLC. L.M.W. is an employee of Adagio Therapeutics Inc. and holds shares in Adagio Therapeutics Inc. L.M.W. is an inventor on a patent describing the ADG anti-SARS-CoV-2 antibodies. W.R.S., D.R.B. and I.A.W. receive research funding from Adagio.

## STRUCTURE DEPOSITIONS

The X-ray coordinates and structure factors have been deposited in the RCSB Protein Data Bank under accession codes: 7U2D and 7U2E.

---

**Supplementary Figure 1.**
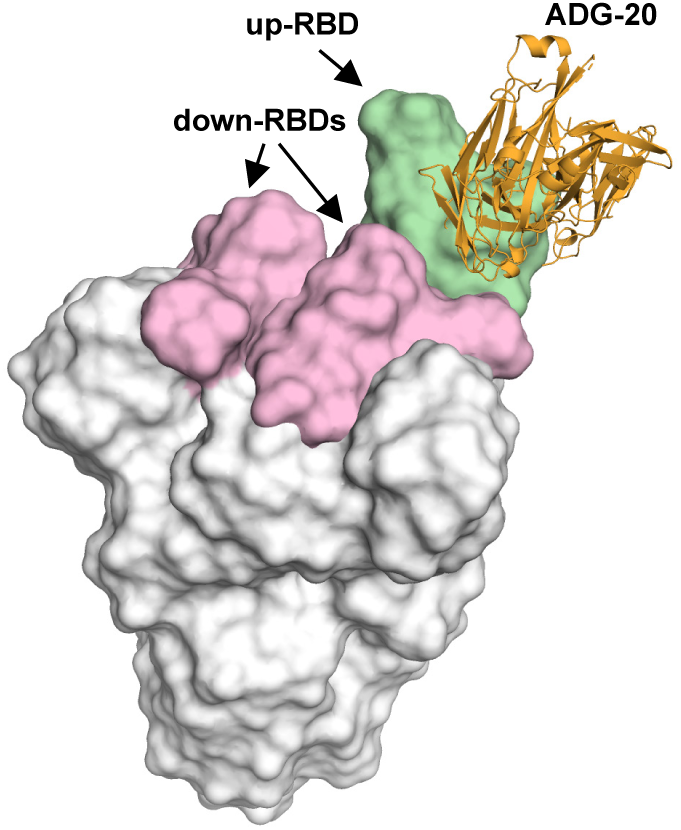
Modelling the binding of ADG-20 on the homotrimeric spike (S) protein. The SARS-CoV-2 S trimer is shown with one RBD in the up conformation (green) and two RBDs in the down conformation (pink). PDB 7ND9 was used in the modeling (*1*). The complete epitope of ADG-20 is accessible only when the RBD is in the up, but not in the down, conformation.

**Supplementary Figure 2.**
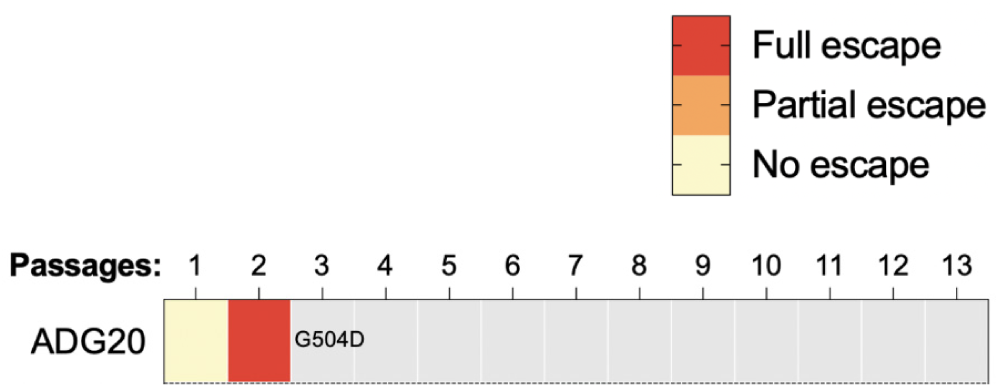
An escape mutant of authentic SARS-CoV-2 against ADG-20. An in vitro SARS-CoV-2 G504D mutant escapes neutralization by ADG-20.

**Supplementary Figure 3.**
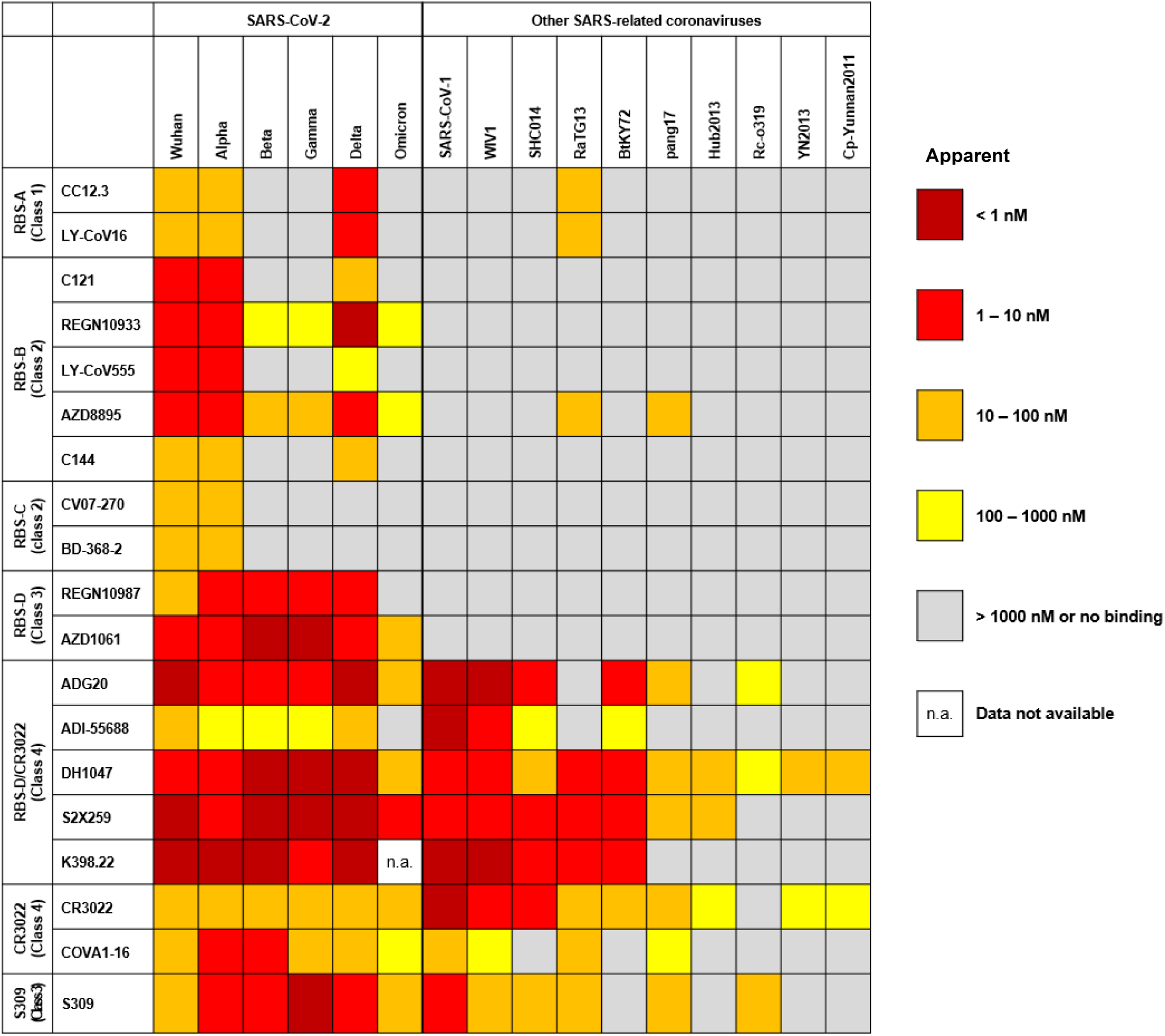
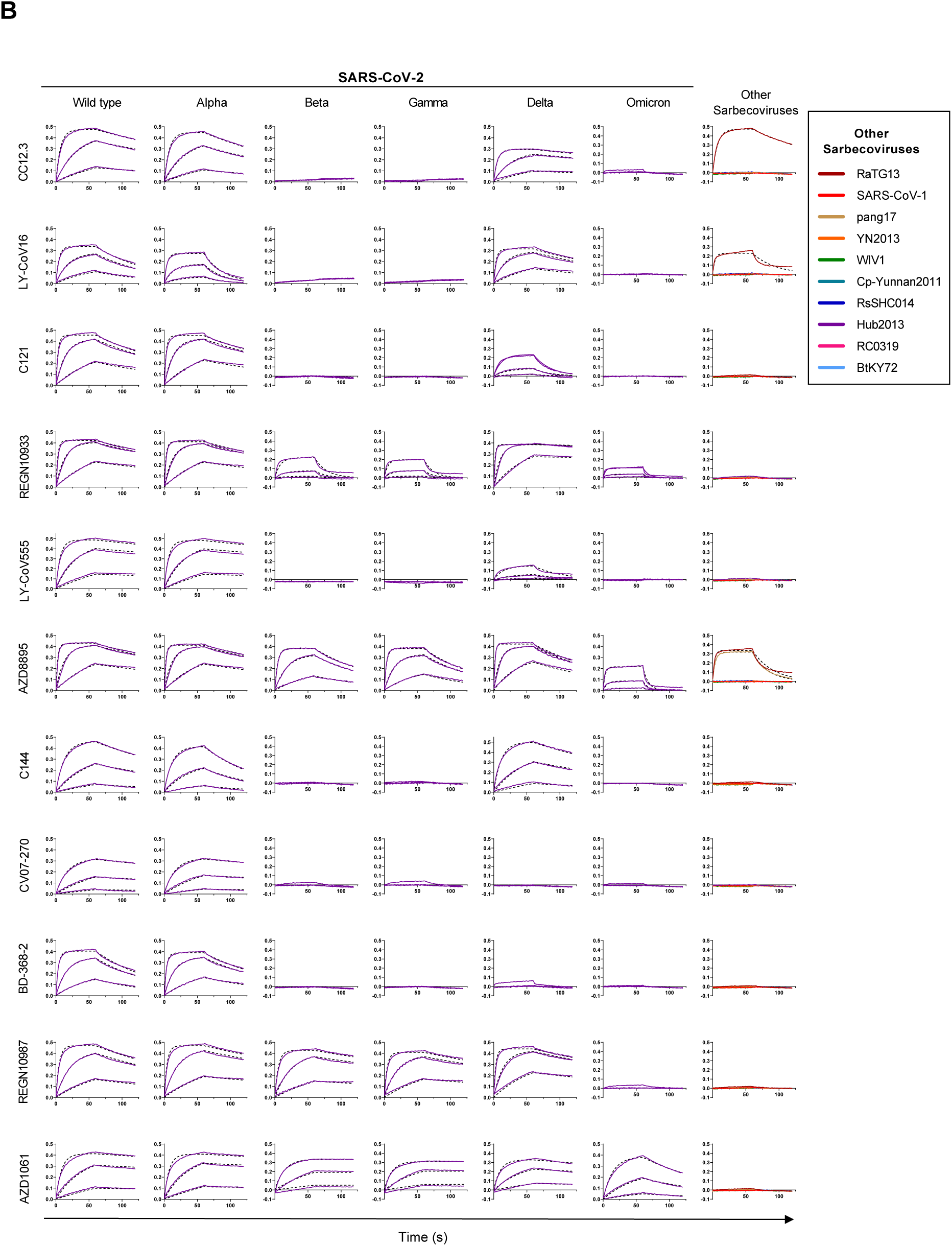
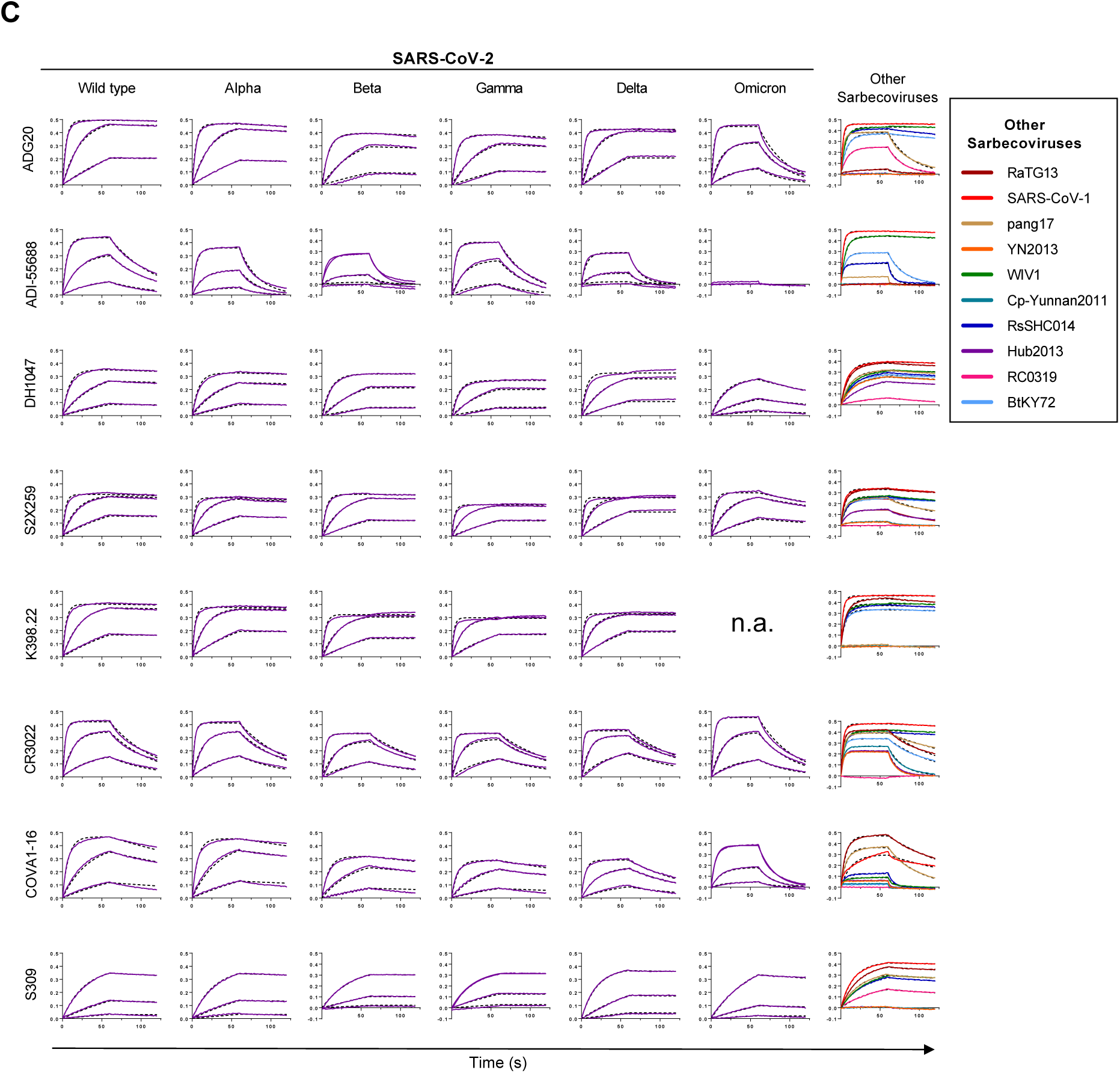
Binding affinity of antibodies with coronavirus spike RBDs. **(A)** BioLayer interferometry (BLI) was used to assess the apparent K_D_, where the IgGs were captured on the sensors and the RBD proteins flowed over. Epitope classification of the RBD antibodies that we previously defined (*2*) is shown in the left column, with the approximate corresponding classes defined in (*3*) shown in brackets. n.a. not available. **(B-C)** Sensorgrams for binding of IgGs and RBDs. The y-axis represents the response against different concentrations of RBD (x-axis in time, seconds). Solid lines represent the response curves and dashed lines represent the 1:1 binding model. The IgGs were loaded onto anti-human IgG Fc (AHC) biosensors and interacted with 5-fold gradient dilution (500 nM – 20 nM) of SARS-CoV-2 RBDs, and 500 nM of RBDs of other sarbecoviruses.

**Supplementary Figure 4.**
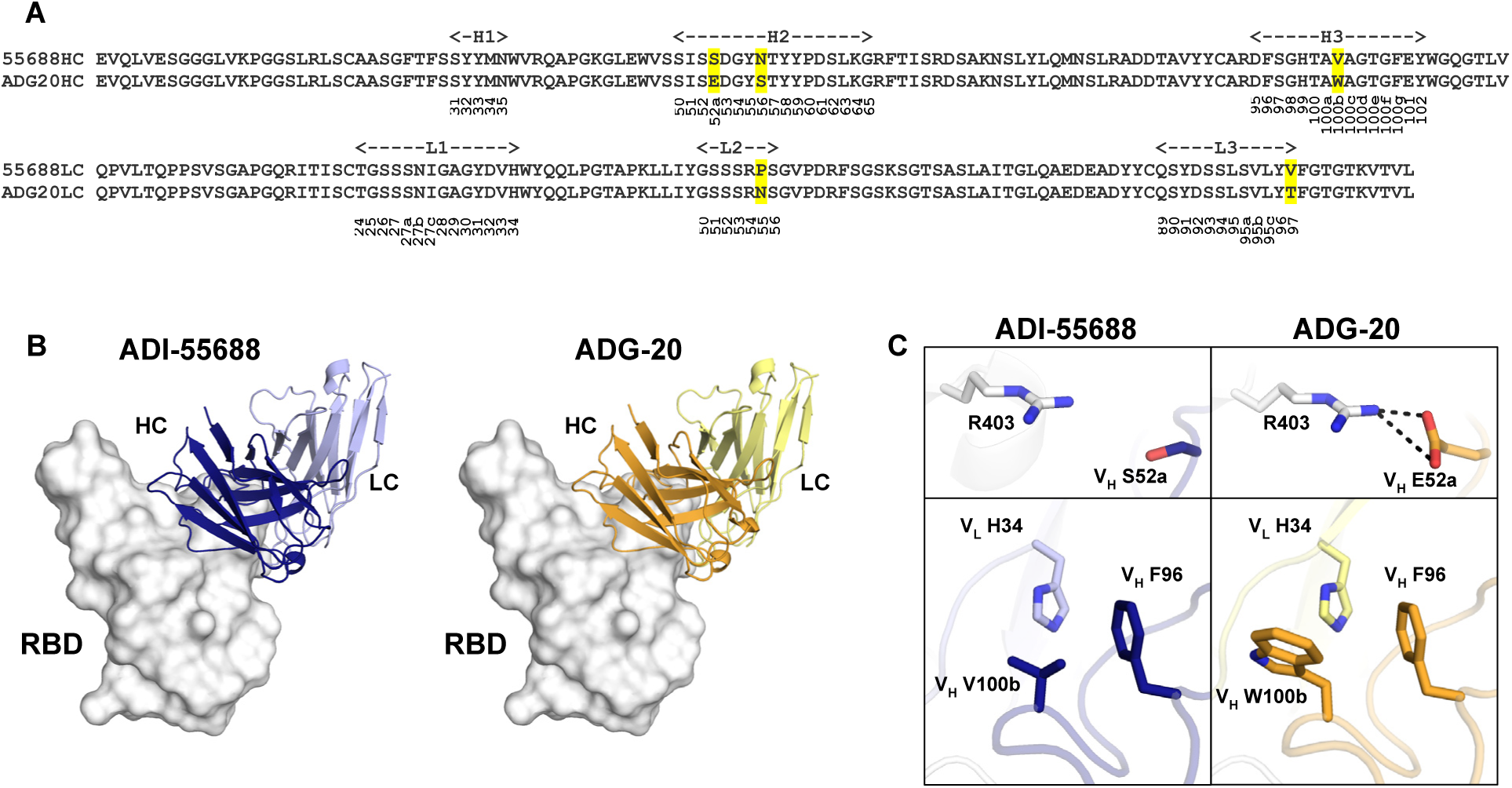
Comparison between ADG-20 and its parent antibody ADI-55688. **(A)** Sequence alignment between ADG-20 and its parent antibody ADI-55688. Different residues are highlighted in yellow. Kabat numbering is shown under the CDR sequences. **(B)** Structural comparison between ADI-55688 and ADG-20. Both antibodies target the same epitope on SARS-CoV-2 RBD (white) in an identical binding mode. **(C)** Detailed interactions of the SARS-CoV-2 RBD with ADI-55688 and ADG-20. In **B** and **C**, heavy and light chains of ADI-55688 are shown in dark and light blue, and those of ADG-20 in orange and yellow. The RBD surface is shown in white. Hydrogen bonds are represented by black dashed lines.

**Supplementary Figure 5.**
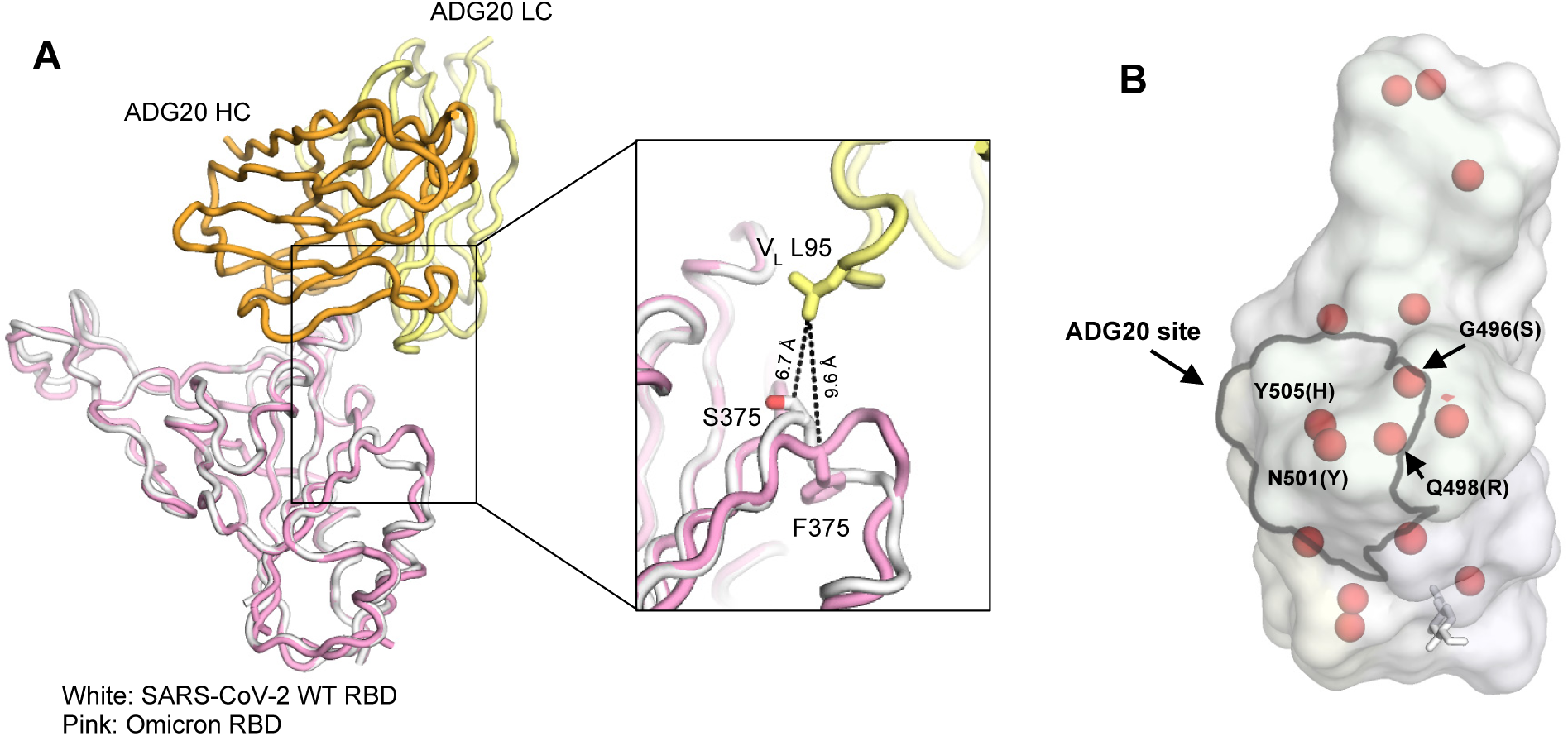
Omicron mutations in the ADG20 epitope. **(A)** Structural comparison between the ADG20-bound wild-type SARS-CoV RBD and Omicron RBD. Crystal structure of ADG20 (heavy chain: orange, light chain: yellow) in complex with wild-type SARS-CoV-2 RBD (white) is from this study (PDB 7U2D). The superimposed Omicron RBD (pink) is extracted from a previous structure in complex with two other Fabs (PDB: 7QNW) (*4*). **(B)** The RBD is represented by a transparent white surface. Mutated residues in the Omicron variant are shown as red spheres. The ADG20 epitope is highlighted by black solid lines. Four residues in the ADG20 epitope are mutated in the Omicron variant: G496S, Q498R, N501Y, and Y505H.

**Supplementary Figure 6.**
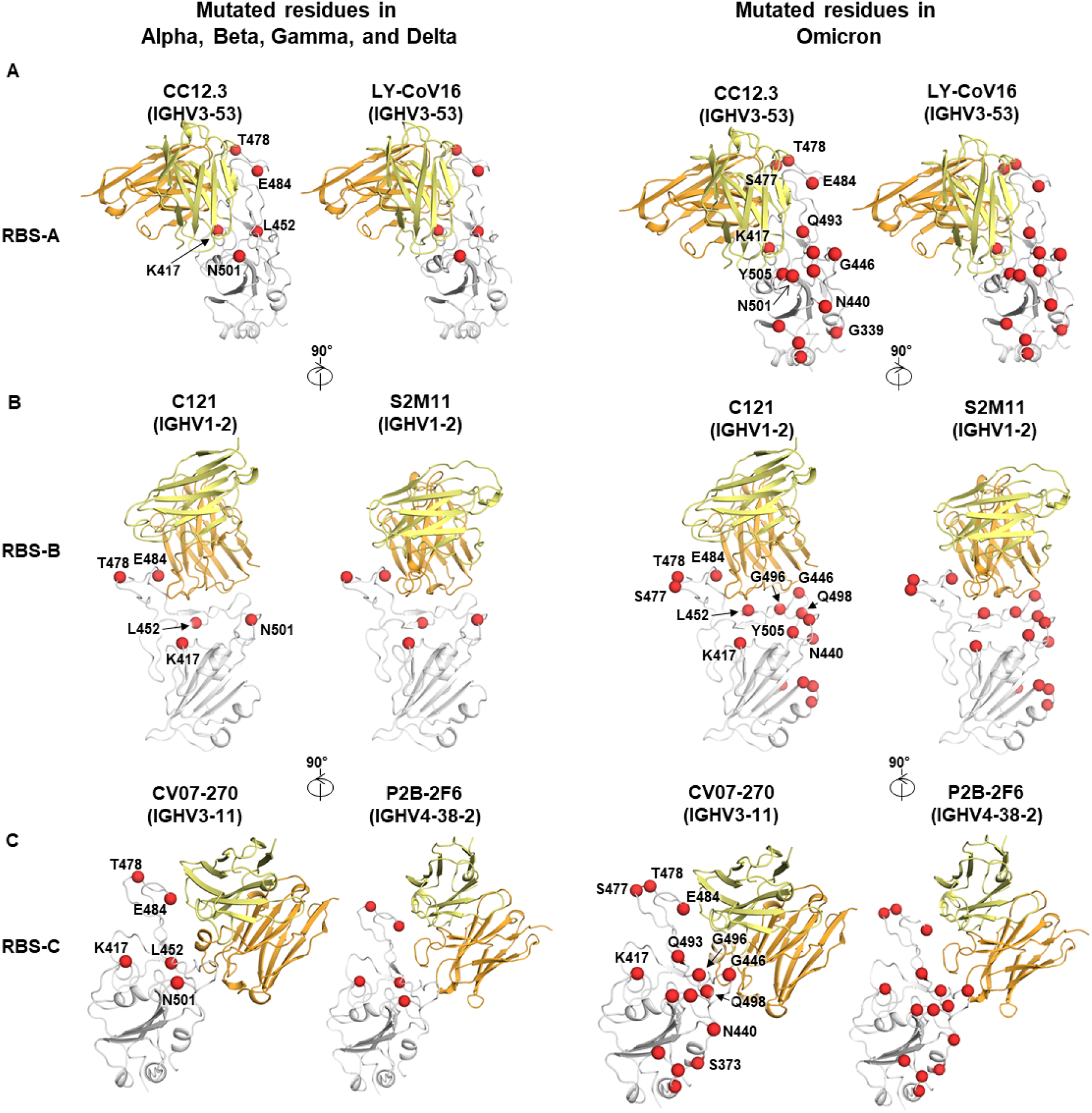
Structures of representative antibodies targeting epitopes RBS-A, B, and C. SARS-CoV-2 RBD is represented by white cartoon with mutated residues in VOCs shown as red spheres, where VOCs Alpha, Beta, Gamma, and Delta are shown in the left panels and Omicron in the right panels. Structures are shown in a same view in each panel. Labelling of some residues are omitted in the right panels for clarity. Heavy and light chains of the bound antibodies are shown in orange and yellow, respectively. Only the variable domains of the antibodies are shown. Structures used for the representative antibodies: CC12.3 (PDB: 6XC4), LY-CoV16/CB6 (7C01), C121 (7K8X), S2M11 (7K43), CV07-270 (6XKP), and P2B-2F6 (7BWJ).

**Supplementary Figure 7.**
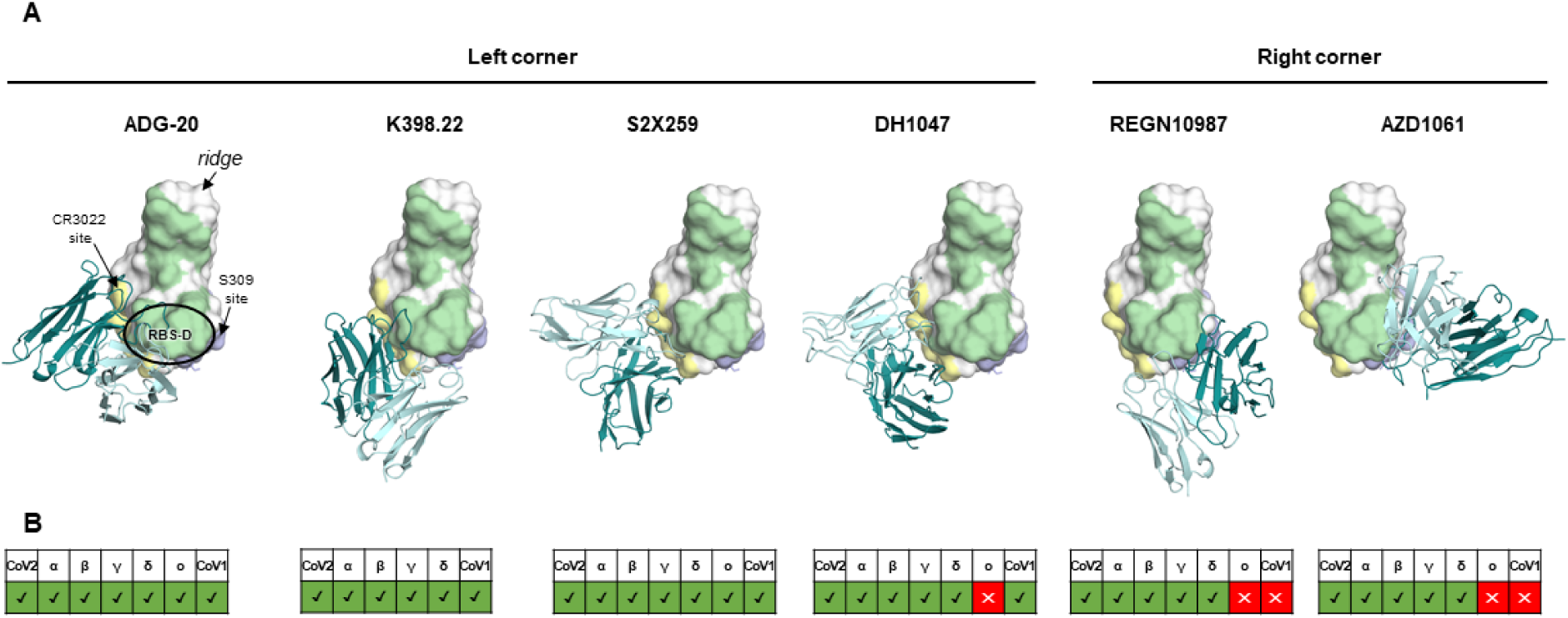
Structural and functional comparison of antibodies targeting opposite corners of RBS-D. **(A)** Structures of RBS-D antibodies. SARS-CoV-2 RBD is shown as white surface with the RBS, CR3022 site and S309 site in green, yellow, and blue, respectively. Heavy and light chains of antibodies are shown as dark and light teal cartoons, respectively. The RBS-D region is highlighted with a black circle in the ADG-20 panel. All panels are in the same view. Structures used in this figure: ADG-20 (this study, PDB 7U2D), S2X259 (PDB 7RAL) (*5*), DH1047 (PDB 7LD1) (*6*), K398.22 (*7*), REGN10987 (PDB 6XDG) (*8*), AZD1061 (PDB 7L7E) (*9*). **(B)** Neutralization of each antibody against SARS-CoV-2 and VOCs, as well as SARS-CoV-1. Each table corresponds to the complex above in panel (A). Antibodies with detectable neutralization are shown in a green “✓” while a red “✕” represents no neutralization. Omicron data for K398.22 are not available.

**Supplementary Table 1.** X-ray data collection and refinement statistics.

| **Data collection** | ADG-20 + SARS-CoV-2 RBD | ADI-55688 + SARS-CoV-2 RBD |
| --- | --- | --- |
| Beamline | APS23ID-B | SSRL12-1 |
| Wavelength (Å) | 1.0332 | 0.9795 |
| Space group | P 4<sub>1</sub> | P 4<sub>1</sub> |
| Unit cell parameters |  |  |
| a, b, c (Å) | 101.1, 101.1, 80.6 | 101.0, 101.0, 79.9 |
| α, β, γ (°) | 90, 90, 90 | 90, 90, 90 |
| Resolution (Å)<sup>a</sup> | 50.0-2.75 (2.81-2.75) | 50.0-2.85 (2.90-2.85) |
| Unique reflections<sup>a</sup> | 20,542 (2,064) | 18,826 (1,811) |
| Redundancy<sup>a</sup> | 11.2 (6.5) | 8.9 (7.3) |
| Completeness (%)<sup>a</sup> | 97.6 (98.1) | 100.0 (99.9) |
| <I/σ<sub>I</sub>><sup>a</sup> | 19.9 (1.0) | 15.1 (1.0) |
| R<sub>sym</sub><sup>b</sup> (%)<sup>a</sup> | 11.4 (98.9) | 14.5 (>100) |
| R<sub>pim</sub><sup>b</sup> (%)<sup>a</sup> | 3.4 (38.3) | 5.1 (51.3) |
| CC<sub>1/2</sub><sup>c</sup> (%)<sup>a</sup> | 99.7 (69.3) | 98.9 (59.0) |
| **Refinement statistics** |  |  |
| Resolution (Å) | 35.7-2.75 | 39.3-2.85 |
| Reflections (work) | 20,538 | 18,820 |
| Reflections (test) | 1,025 | 1,882 |
| R<sub>cryst</sub><sup>d</sup> / R<sub>free</sub><sup>e</sup> (%) | 21.9/25.8 | 25.4/28.4 |
| No. of atoms | 4,873 | 4,837 |
| RBD | 1,561 | 1,561 |
| Fab | 3,240 | 3,231 |
| Ligands<sup>f</sup> | 38 | 45 |
| Solvent | 34 | 0 |
| Average B-values (Å<sup>2</sup>) | 72 | 66 |
| RBD | 69 | 67 |
| Fab | 73 | 65 |
| Ligands | 119 | 68 |
| Solvent | 56 | N/A |
| Wilson B-value (Å<sup>2</sup>) | 72 | 70 |
| **RMSD from ideal geometry** |  |  |
| Bond length (Å) | 0.002 | 0.002 |
| Bond angle (°) | 0.57 | 0.57 |
| **Ramachandran statistics (%)<sup>g</sup>** |  |  |
| Favored | 96.1 | 96.1 |
| Outliers | 0.16 | 0.00 |
| **PDB code** | 7U2D | 7U2E |

<sup>a</sup> Numbers in parentheses refer to the highest resolution shell.
<sup>b</sup> $R_{sym} = \sum_{hkl} \sum_i |I_{hkl,i} - \langle I_{hkl} \rangle| / \sum_{hkl} \sum_i I_{hkl,i}$ and $R_{pim} = \sum_{hkl} (1/(n-1))^{1/2} \sum_i |I_{hkl,i} - \langle I_{hkl} \rangle| / \sum_{hkl} \sum_i I_{hkl,i}$ , where $I_{hkl,i}$ is the scaled intensity of the $i^{th}$ measurement of reflection $h, k, l$ , $\langle I_{hkl} \rangle$ is the average intensity for that reflection, and $n$ is the redundancy.
<sup>c</sup> CC<sub>1/2</sub> = Pearson correlation coefficient between two random half datasets.
<sup>d</sup> $R_{cryst} = \sum_{hkl} |F_o - F_c| / \sum_{hkl} |F_o| \times 100$ , where $F_o$ and $F_c$ are the observed and calculated structure factors, respectively.
<sup>e</sup> $R_{free}$ was calculated as for $R_{cryst}$ , but on a test set comprising 5% or 10% of the data excluded from refinement.
<sup>f</sup> Bound ligands are SO<sub>4</sub> and citrate.
<sup>g</sup> From MolProbity (10).

**Supplementary Table 2.** Hydrogen bonds and salt bridges identified at the antibody-RBD interface using the PISA program.

| ADG-20 | Distance [Å] | SARS-CoV-2 RBD |
| --- | --- | --- |
| **Hydrogen bonds** |  |  |
| VH:SER56[OG] | 3.8 | ASP405[O] |
| VH:TYR33[OH] | 2.7 | ASP405[OD2] |
| VH:THR100E[OG1] | 3.3 | THR500[O] |
| VH:GLU52A[OE2] | 3.7 | ASN501[OD1] |
| VH:THR100[O] | 3.0 | GLN498[NE2] |
| VH:HIS99[O] | 3.2 | GLN498[NE2] |
| VH:ALA100A[O] | 2.5 | THR500[OG1] |
| VH:ALA100A[O] | 3.7 | ASN501[N] |
| VH:THR100[O] | 3.5 | ASN501[ND2] |
| VH:THR100E[OG1] | 3.8 | GLY502[N] |
| VH:ALA100C[O] | 2.8 | GLY502[N] |
| VH:GLU52A[OE2] | 2.8 | TYR505[OH] |
| **Salt bridges** |  |  |
| VH:GLU52A[OE1] | 3.8 | ARG403[NH2] |
| VH:GLU52A[OE2] | 2.5 | ARG403[NH2] |

| ADI-55688 | Distance [Å] | SARS-CoV-2 RBD |
| --- | --- | --- |
| **Hydrogen bonds** |  |  |
| VH:TYR33[OH] | 2.8 | ASP405[OD2] |
| VH:THR100E[OG1] | 3.3 | THR500[O] |
| VH:ALA100A[O] | 2.4 | THR500[OG1] |
| VH:ALA100A[O] | 3.7 | ASN501[N] |
| VH:THR100[O] | 3.5 | ASN501[ND2] |
| VH:ALA100A[O] | 2.9 | ASN501[ND2] |
| VH:ALA100C[O] | 2.6 | GLY502[N] |
| VH:TYR33[OH] | 2.8 | ASP405[OD2] |
| VH:THR100E[OG1] | 3.3 | THR500[O] |
| L:TYR31[OH] | 2.7 | GLN506[NE2] |
| L:TYR91[OH] | 3.0 | VAL503[N] |

**Supplementary Table 3.** Putative germline genes and CDR H3 sequences of antibodies targeting the RBS-D/CR3022 epitope.

|  | **IGHV** | **IGK(L)V** | **CDR H3 sequence** | **Reference** |
| --- | --- | --- | --- | --- |
| **ADG-20** | IGHV3-21 | IGLV1-40 | ARDFSGHTAWAGTGFEY | (11), this study |
| **DH1047** | IGHV1-46 | IGKV4-1 | ARDVRVDDSWSGYDLLSGGTYFDY | (6, 12) |
| **S2X259** | IGHV1-69 | IGLV1-40 | ARGFNGNYYGWGDDDAFDI | (5) |
| **K398.22** | Macaque<br>IGHV3-73 | Macaque<br>IGLV2-32 | TRVSIFGQFIVATYFDY | (7) |

